# Analysis of lncRNA, miRNA, mRNA-associated ceRNA networks in Autism

**DOI:** 10.1101/2023.05.14.540737

**Authors:** Asal Tabatabaei Azad, Mohammad-Reza Mehrabi, Yasamin Zarinzad, Atefeh Noori, Hossein Nemati, Amir Shahbazi, Parisa Badameh, Seyed Roham Mohammadnezad kenari, Parham Arabzadeh, Amir-Reza Javanmard

## Abstract

Autism is a multifactorial behavioral disease, which is caused by different environmental and genetic alterations. In this disease, various molecular pathways such as inflammation and cell-cell connections are affected. In this study, by analyzing RNA-seq data and bioinformatics approaches, candidate genes were selected for further analysis, and the expression of candidate genes in the blood samples of autistic people was compared to normal. Finally, the candidate genes showed potential as biological markers for prognosis in autism. Through the analysis of these candidate genes, the researchers were able to identify changes in the expression of certain genes that were associated with the disease. By understanding how these genes are involved in the disease, researchers can develop better treatments and diagnose the disease earlier.

## Introduction

Autism spectrum disorder (ASD) is a neurodevelopmental disorder and a complex syndrome consisting of a wide range of symptoms caused by abnormalities in the structure and function of the brain. This condition is characterized by difficulties communicating and social skills, delayed speech and motility, abnormal reactions to sound and other stimuli, repetitive behaviors and interests, and digestive problems. Autism is inherited in 80% of cases but environmental factors, random mutations and epigenetics can also play a role in its occurrence[1-3].

The prevalence of autism around the world has increased in recent years due to advances in screening and diagnostic methods. In the United States in 2022, the prevalence of autism spectrum disorders was about 1 in 44 children. Moreover, it was similar between races and there was no difference between black and white children. ASD is potentially affected by gender, with boys being about 4 times more likely to be diagnosed than girls.

Autism symptoms usually appear in the first three years of life and it can be diagnosed at the age of 18 months or even earlier, although the diagnosis is more reliable at 2 years of age[4]. However, some people are not diagnosed until adolescence. Early detection of autism is a vital step, for it has a significant impact on improving symptoms and raising the life quality of ASD children.

. According to science, ASD gender-dependence stems from the fact that women have two x chromosomes and alterations to one chromosome can be compensated by another healthy chromosome.But for men, having only one x chromosome, defects on the x chromosome can lead to an unhealthy phenotype. For instance, a defect in the *NLGN4* gene, which acts to create and maintain synaptic structures and is located on two sex chromosomes (one on the x chromosome and one on the y chromosome), has been shown to be involved in autism. *NLGN4Y* protein does not have the required function due to having a different single amino acid and the main function of this protein is related to *NLGN4X* protein. The region around this amino acid in the *NLGN4X* gene is a mutation-sensitive region, and different autism-associated variants have been seen in people with autism[5].

Genomic studies since 2005 have led to the identification of numerous CNVs, SNVs and SNPs, with CNVs accounting for about 15% and SNVs for 7% of autism cases. However, their association with autism is limited to a small number of people and has a very low prevalence in society[6, 7]. In addition, more than 1,000 autism-related genes are candidates in case-control studies. Most of these genes encode proteins involved in synaptic structure and function, and the others have functions in cell proliferation, chromatin rearrangement, and transcriptional regulation (released 20 July 2022, gene.sfari.org).

In humans, almost 80% of the genome is transcribed, with only 2% of these transcripts encoding proteins and the rest including non-coding RNA[8]. Two main groups of ncRNAs include microRNAs (miRNA) and long noncoding RNAs (lncRNAs), which have a regulatory role in various cellular processes.Changes in them are related to the occurrence of diseases and abnormalities[9].

To date, a large number of ncRNAs are known to be associated with neurodevelopment. Changes in their expression level are related to neurodevelopmental diseases, including autism[10]. Analysis of circulating serum samples from children with autism has led to the identification of a number of differentially expressed miRNAs[11]. The interaction between these miRNAs and the critical ASD genes indicates a potential role for the ncRNAs in the molecular pathways involved in ASD. Moreover, examining prefrontal cortex and cerebellum tissues of Postmortem brain, identified more than 200 autism-related lncRNAs with regions containing genes associated with neurodevelopmental disease[12]. LncRNAs can affect the regulation of gene expression through sponging miRNAs[13]. The study of cross-talk between miRNA-lncRNA-mRNA as a ceRNA network analysis has provided a way to identify biomarkers and molecular pathways associated with autism.

In this study we aimed to identify and screen potential biomarkers for ASD. We also studied the interactions between differentially expressed RNAs in the autistic brain through the construction and analysis of a CERNA network.. Based on the connections between DEmRNAs, DEmiRNAs and DElncRNAs we were able to identify a biomarker for ASD.

## METHODS

### Differential expression of lncRNAs, mRNAs, and microRNA

In order to investigate and find differentially expressed RNA, including mRNA, microRNA and LncRNA in autism, the raw data of sequencing (*SRP174367*) was taken from the SRA database (http://www.ncbi.nlm.nih.gov/sra). The selection of raw data was done based on a search for the word autism. Information files related to phenotype and expression level were extracted from primary data. Genes were annotated using HTSeq-Count and then 16 numbers of autism samples and 14 healthy samples were compared in terms of expression levels. Using the R package edgeR with the threshold of P-value < 0.05 and |log2-fold change (FC)| > 2 dRNAs were determined and then used for further analysis.

### lncRNA-miRNA-mRNA related network

Mircode database (Version 11; http://www.mircode.org/mircode/, accessed on 25 June 2022), targetscan (Version 8.0; http://www.mircode.org/mircode/, accessed on 25 June 2022) and starbase were used to investigate the interaction between demRNA, demiRNA and lncRNA and to predict the direct relationship between them. The mircode database makes it possible to check the interactions and binding sites of miRNAs among the complete GENCODE annotated transcriptome. Through it, the binding of Mir lncRNAs can also be checked [8]. The interactions between Mir and mRNA were explored using the Mirbase, TargetScan and Starbase databases. Starbase explores Mir targets with data obtained from CLIP-Seq and Degradome-Seq[9]. The possible targets for miR were predicted based on the complementary relationship and connection between the miR seed region and the 3UTR region of mRNA.

### The ceRNA regulatory network

The interactive relations between the differentially expressed miRNAs, mRNAs lncRNAs were derived from pre-named databases. Then the predicted miRNA-lncRNA interactions and miRNA-mRNA interactions were visualized as a novel ceRNA network related to autism using Cytoscape software.

### Enrichment analysis

To classify the genes based on their function either in molecular function(MF), biological processes(BP) and cellular components(CC) the Gene Ontology(GO) analysis and to map genes in pathways Kyoto Encyclopedia of Genes and Genomes (KEGG) enrichment analysis were made by the use of R.

## Results

### Differential expression analysis of lncRNAs, mRNAs, and miRNAs from public SRA data

Differential expression analysis led to 499 DEGs 150 DEMs and 550 DELs, from which 225 mRNA 75 miRNA 260 lncRNA were upregulated and 274 mRNA 75 miRNA 290 lncRNA were downregulated (table1&2). All were evaluated with the criteria of logfc< 0.5 or >2 and pv <0.05. expression level of differentially expressed mRNA and lncRNAs is shown in heatmap and volcano plot .(fig1)

**Fig1).**
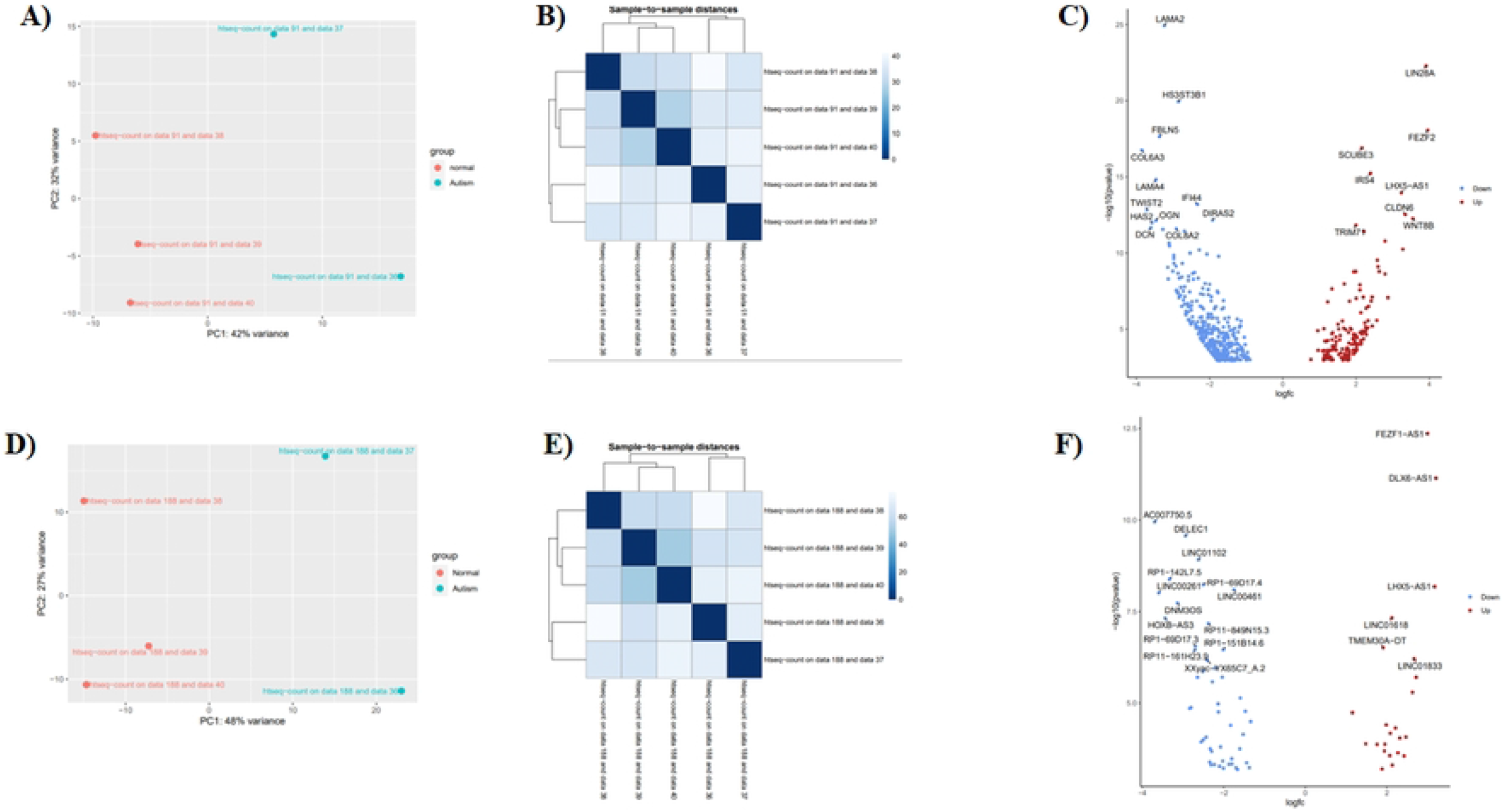
PCA plot and heat map diagram for dysregulated genes. A) The PCA diagram shows that the autism samples are completely normal. B) The Heatmap diagram shows that the autism samples are completely normal. C) The volcano plot shows the distribution of dysregulated mRNAs. D) The PCA diagram shows that the autism samples are completely normal. E) The Heatmap diagram shows that the autism samples are completely normal. F) The volcano plot shows the distribution of dysregulated LncRNAs.

### Prediction and construction of the lncRNAs-miRNAs-mRNAs network

A lncRNA–miRNA–mRNA regulatory network was constructed based on the links between miRNA and mRNA and also between miRNA and lncRNA. Possible interactions were predicted using starbase, micode and targetscan. A competitive endogenous RNA network is built to describe lncRNA functions in competing with miRNAs in binding to mirs and also sponging them in order to regulating target genes expression (fig2).

**Fig2).**
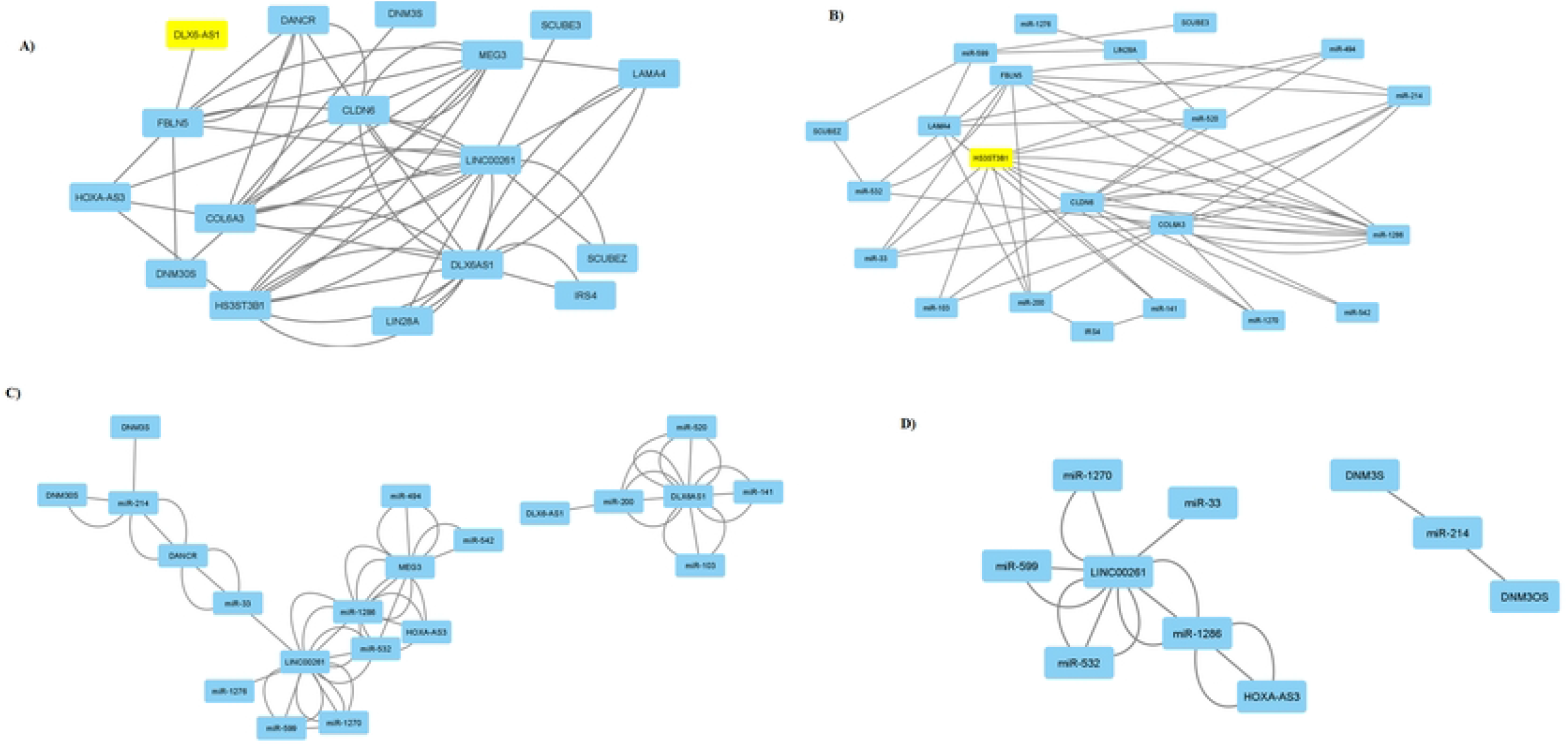
The lncRNA-miRNA-mRNA ceRNA network analysis in Autism. A&B) The lncRNA-miRNA-mRNA ceRNA network analysis in Autism. C&D) The main ceRNA networks in Autism.

### Functional and pathway enrichment analysis

By using enrichers, GO analysis containing Cellular Components (CC), Biological Functions (BF) and Molecular Processes (MP) and pathway analysis was performed to evaluate DEGs functions.The KEGG database was also examined to indicate the involvement of DELs in different pathways. Accordingly, 14 MF, BP, CC and pathways were associated with DEGs (p<0.05). Top GO analysis and KEGG pathways are shown in (fig3).

**Fig3).**
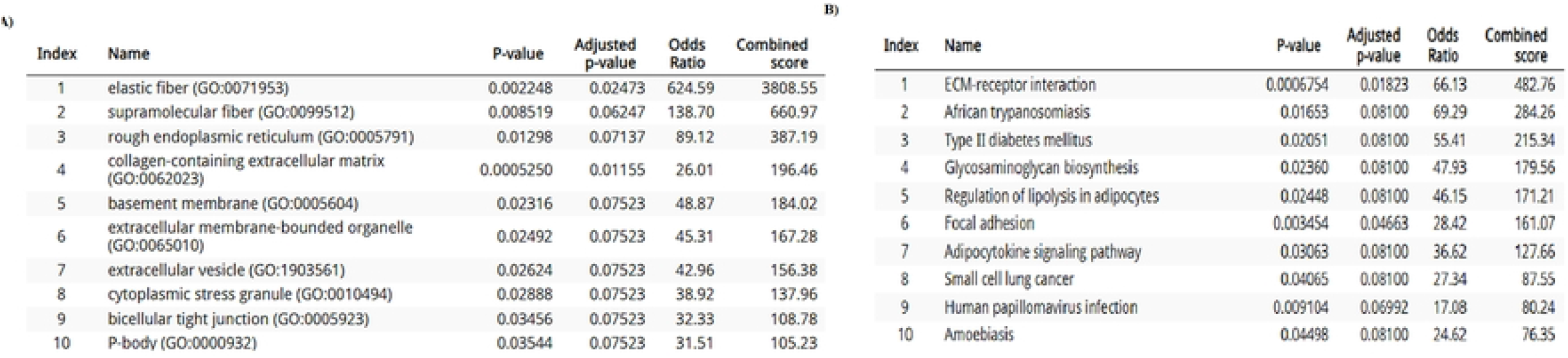
Predicted cellular and molecular pathways involved in autism. A&B) KEGG analysis predicted pathways involved in autism.

## Discussion

Autism spectrum disorder is a type of neurodevelopmental spectrum disorder that is characterized by abnormal communication and verbal behaviors[14]. The symptoms of this disorder usually appear before the age of 1 to 3 years and its cause is still unknown, but it is believed that genetic factors and the environment are involved in it. The social, economic, lifestyle, and education of parents do not affect this disorder[15]. Because of the heterogeneity among ASD patients, the diagnosis of ASD depends on the assessment of the patient’s behavior. Therefore, identification Specific molecular markers may help to understand the pathophysiological mechanism of ASD, thereby increasing the chances of correct diagnosis and improving therapeutic interventions [16]. Many studies have shown that females with ASD show different characteristics than males with ASD. In addition, clinical studies have also reported that according to the different characteristics of ASD and camouflage, females with ASD are diagnosed much later than males with ASD[17]. Many genes are affected by risk mutations In ASD encoded proteins for synaptic, transcriptional and Chromatin remodeling pathways, which involve voltage ion channels regulating the release of action potentials, Acceleration and excitability-transcription coupling as well as histone modifying and chromatin remodeling enzymes[18]. In recent studies, it has been found that the molecular pathways of the inflammatory response[19] and PI3K[20], MAP Kinase[21], and TGFB play significant roles in the development of autism[3]. in the process of brain development, PI3K involves in different cellular functions, including cell migration, propagation, and axonal guidance[22]. Inactivation of PTEN (an upstream negative regulator of PI3K-AKT/mTOR pathway) causes upregulation of PI3K-AKT/mTOR pathway that leading to axonal dysregulation, megalocephaly, alteration of neuron size, protein synthesis, cerebral cell proliferation, and neuronal circuit connectivity in different regions of the brain[20]. transforming growth factor beta (TGFB) consists of three closely related isoforms TGFB1-TGFB2-TGFB3. These three isoforms are expressed in several central nervous system CNS cell types including neurons, astrocytes, and microglia[3]. In the past, it has been observed that the expression level of TGFB1 in the plasma of autistic children is significantly reduced[23]. The MAPK pathway regulates neural progenitor biogenesis, learning and memory, and mRNA translation by phosphorylation of TSC2 and eIF4E through MAPK-interacting kinase 1 and 2 (MNK1 and MNK2). Besides, The deletion of *MAPK3* (mitogen-activated protein kinase 3) gene that is located in 16p11.2 locus is related to ASD patients[24]. Moreover, ERK-MAPK signaling pathway in neurons has a pivotal function in regulating of synaptic plasticity[25] the MAPK pathway in patients with ASD syndromic is upregulated[26]. In addition, it has been found that LncRNAs and MicroRNAs, which are ncRNAs[27]. play a significant role in physiological processes[28]. Cellular and also in causing or preventing various diseases including cancers, diabetes, cardiovascular diseases, and nerve diseases[29]. For example, lncRNAGAS5 causes autism by affecting mir-221 on the PI3K signaling pathway[30]. Also in the early stage of brain development, Lin28A mediates increased protein synthesis that is crucial for neural progenitor cell, proliferation and maintenance.

Furthermore, exogenous LIN28A promoted the proliferation of neural progenitor cells in vitro and induced early survival effect on primary cortical neurons, possibly via Upregulation of IGF2[31]. Our bioinformatics analysis revealed a number of lncRNAs, microRNAs, and mRNAs involved in pathways such as inflammation, WNT, TGFB, and PI3K, which are significant pathways in this disease. Such as the genes *LIN28A*[32], *CLDN6*[33], *COL6A3*[34], *FBLN5*[34], and LncRNAs *HOXA-A3, DANCR, LINC00461, LINC00261*, and microRNAs *miR-103, miR-494, miR-520, miR-141, miR-214, miR-33*.

## Conclusion

The construction of ceRNA network introduces Several genes that are regulated by various factors through the influence they have on each other via multiple signaling pathways. As a final conclusion, it can be suggested that the identified genes in this study may serve as novel biomarkers for the diagnosis of autism and may provide insights into how the interaction of environmental factors with the genetic of an individual lead to the development of autism. This could potentially help to create more accurate diagnostic tests that can be used to identify autism in its early stages. Furthermore, this research may open the door to new treatment and prevention strategies for autism, enabling individuals to better manage and live with this condition.

## Declarations

### Ethics approval and consent to participate

This study was approved by the Ethical/Scientific Committee of Tarbiat Modares University (IR.MODARES.REC.1400.122).

### Competing interests

The authors declare that they have no competing interests.

### Funding

There was no funding for this study.

### Author contributions

ASTA,MM,YZ&AN performed the experiments. ARJ designed the experiments and supervised the study. AS,SRK,PA&PB helped with lab works.

## Acknowledgments

Authors are thankful for the help and advice of the 4402-laboratory members at the Department of Genetics, TMU. Tehran-Iran

## Notes

### Competing Interest Statement

Enter: The authors have declared that no competing interests exist.

